# Overexpression of Alpha-1 Antitrypsin Increases the Proliferation of Mesenchymal Stem Cells by Upregulation of Cyclin D1 and is Independent of the Wnt Signaling Pathway

**DOI:** 10.1101/2023.10.28.564526

**Authors:** Bryan Wolf, Prasanth Muralidharan, Michael Lee, Wei Hua, Erica Green, Hongjun Wang, Charlie Strange

## Abstract

Alaph-1 antitrypsin overexpressing mesenchymal stromal/stem cells (AAT-MSCs) showed improved innate properties with a faster proliferation rate when studied for their protective effects in mouse models of diseases. Here, we investigated the potential mechanism(s) by which AAT gene insertion increases MSC proliferation. Human bone marrow-derived primary or immortalized MSCs (iMSCs) or AAT-MSCs (iAAT-MSCs) were used in the study. Cell proliferation was measured by cell counting and cell cycle analysis. Possible pathways involved in the pro-proliferation effect of AAT were investigated by measuring mRNA and protein expression of key cell cycle genes. Interval cell counting showed increased proliferation in AAT-MSCs or iAAT-MSCs compared to their corresponding MSC controls. Cell cycle analysis revealed more cells progressing into the S and G2/M phases in iAAT-MSCs, with a notable increase in the cell cycle protein, Cyclin D1. Moreover, treatment with Cyclin D1 inhibitors showed that the increase in proliferation is due to Cyclin D1 and that the AAT protein is upstream and a positive regulator of Cyclin D1. Furthermore, AAT’s effect on Cyclin D1 is independent of the Wnt signaling pathway as there were no differences in the expression of regulatory proteins, including GSK3β and β-Catenin in iMSC and iAAT-MSCs. In summary, our results indicate that AAT gene insertion in an immortalized MSC cell line increases cell proliferation and growth by increasing Cyclin D1 expression and consequently causing cells to progress through the cell cycle at a significantly faster rate.

## INTRODUCTION

The use of adult stem cells to treat a variety of diseases represents a novel therapeutic approach for many diseases (1). Stem cells are particularly effective in healing because they can proliferate and differentiate into specialized cells in the body. These properties allow them to grow and develop as replacements for cells that have been damaged or have died. The ability of a stem cell to differentiate defines part of its therapeutic potential (1). Pluripotent stem cells can differentiate into specialized cells like neurons or skin cells; however, multipotent stem cells are more limited in their potential differentiation (2). The cells used in this study were mesenchymal stem/stromal cells (MSCs). MSCs are multipotent stem cells that can differentiate into adipocytes, chondrocytes, and many other types of cells. These cells are predominantly found in bone marrow can also be derived from umbilical cords and several other tissues (3, 4). MSCs are unique that they are relatively easy to acquire compared to other stem cells as they can be harvested from adult tissue as opposed to the human embryonic tissue. MSCs are an effective treatment for various injuries and illnesses and can display anti-inflammatory properties (1, 2).

There are various therapeutic benefits of the protein alpha-1-antitrypsin (AAT). AAT is a protease inhibitor produced in the liver protecting the lungs and other organs from neutrophil elastase damage (5). AAT exhibits unique properties such as apoptosis inhibition and promotion of cell proliferation (3, 4, 6, 7). Previous studies have indicated that human AAT-overexpressing mesenchymal stem cells (AAT-MSCs) improved the innate properties of naive MSCs and showed improved therapeutic effects in animal models of type 1 diabetes and graft vs host disease (6–8) (9).

Glycogen Synthase Kinase 3 Beta (GSK3β) is a protein that controls the rate and progression of cellular proliferation. The protein does this by acting as a negative regulator of other major cell cycle proteins such as Cyclin D1 (Cyc D1) (10). Cyclin D1 plays a crucial role in positively regulating the expression of cyclin D-dependent kinases that, when activated, will catalyze DNA synthesis and mitosis (11). Cyclin D1 has been linked to some forms of cancer from the lack of regulation of Cyclin D dependent kinases (12). GSK3β kinase works to phosphorylate and control the expression of mitotic proteins like Cyclin D1 regulating cell growth and when GSK3β is phosphorylated it allows Cyclin D1 to be expressed. This activation of Cyclin D1 causes cells to initiate DNA replication and progress into the DNA synthesis (S phase) part of their cell cycle (10).

The Wnt/β-Catenin signaling pathway also has an important role in cell growth and cell development. One of the many functions of the pathway is the regulation of cell proliferation (13–15). One result of the Wnt pathway is the expression and translocation of β-catenin into the nucleus, leading to upregulation of the expression of several genes related to cell proliferation including Cyclin D1 (13). GSK3β is also involved in this pathway as it regulates upstream the activation of β-Catenin after stimulation from Wnt transmembrane receptors (13). When GSK3β is inhibited, it allows the pathway to continue and stimulate cellular growth. Additionally, GSK3β can regulate and degrade β-catenin and control Cyclin D1 expression independent of the Wnt pathway (13). The goal of this study is to investigate the mechanisms of how AAT overexpression in immortalized MSCs led to increased proliferation by analyzing the potential involvement of Cyclin D1 and other proteins involved in cell proliferation.

## RESULTS

### 1. iAAT-MSCs showed higher proliferation compared to control iMSCs

To compare cell growth rate, bone marrow-derived iMSCs and iAAT-MSCs at the same passage (P7-P9) were seeded at the same density, and cell proliferation rate was measured by counting cell numbers at day (D) 1, 3, 5 and 7 after seeding. Both iMSCs and iAAT-MSCs were GFP^+^ cells that showed typic features of MSCs (**Figure 1A**). Our data show that iAAT-MSCs showed faster growth compared to corresponding control iMSCs at day 5 (**Figure 1B**). By day 7 the difference disappeared, likely due to over confluence.

**Figure 1:**
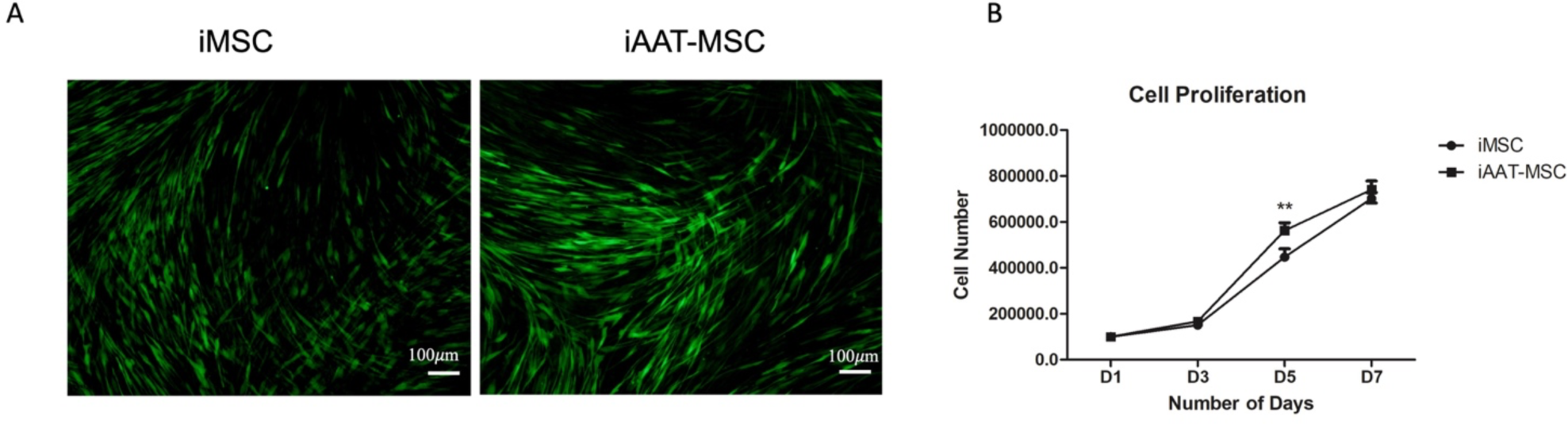
iAAT-MSCs showed significant increases in cell proliferation compared to control iMSCs. (A) Sample immunofluorescent image of GFP^+^ iMSCs and iAAT-MSCs. Magnification x10. (B) Cell number at 1, 3, 5, and 7 days post-seeding, with iAAT-MSCs having a higher cell number than normal iMSCs at day 5. **, *Student’s t* test, p<0.01.

### 2. Cell cycle analysis showed a difference between iAAT-MSCs and iMSCs

To confirm the difference in cell proliferation, a cell cycle analysis was conducted. As shown in **Figure 2A**, there were significantly fewer iAAT-MSC cells in the G0/G1 phase than their iMSC counterparts. Accordingly, there was an increase in the peak that corresponds to the S and the G2/M phases in iAAT-MSCs compared to the iMSCs (p < 0.05, **Figure 2B**). These results suggest that iAAT-MSCs exhibit a faster mitosis capacity due to a significant increase in the numbers of cells that progressed from G0/G1 to S and G2/M compared with iMSCs.

**Figure 2:**
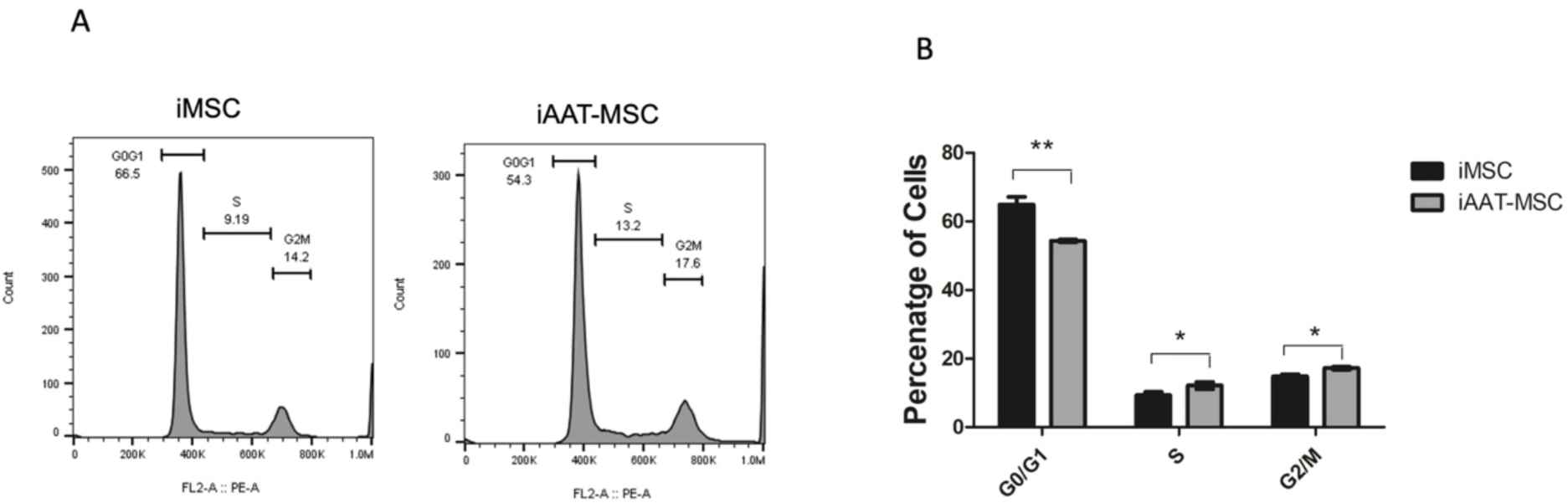
Cell cycle analysis showed increases in S and G2/M Phases in iAAT-MSCs compared to iMSCs. (A). Representative histogram shows cell cycle of iMSCs and iAAT-MSCs analyzed by flow cytometry (B) Mean percentage of cells in the G0/G1, S, and G2/M phase in iMSCs and iAAT-MSCs from 3 separate experiments. *p<0.05, ** p<0.01, ANOVA.

### 3. Upregulation of Cyclin D1 in iAAT-MSCs compared to iMSCs

We next assessed the potential causes of this increase in proliferation in iAAT-MSCs. We conducted a single cell RNAseq analysis to define possible genes involved in the increased proliferation of AAT-MSCs vs MSCs, and an upregulation of the Cyclin D1 gene in the iAAT-MSCs was observed (Wang, unpublished data). We next used qPCR to confirm Cyclin D1 gene expression. An increase in Cyclin D1 mRNA expression was observed in iAAT-MSCs compared to control iMSCs (p<0.05) (**Figure 3A**). This increase in gene expression translated to protein levels as both total and phosphorylated Cyclin D1 (p-Cyclin D1) were higher in the iAAT-MSCs compared to iMSCs (**Figure 3B&C**). However, the p-Cyclin D1/ total Cyclin D1 ratios were similar between iMSCs and iAAT-MSCs, suggesting an increase in absolute Cyclin D1 (**Figure 3D**). The rise in Cyclin D1 protein expression in the iAAT-MSCs is also demonstrated qualitatively by the increase in fluorescence density in immunofluorescent staining of Cyclin D iAAT-MSCs vs. iMSCs (**Figure 3E**).

**Figure 3:**
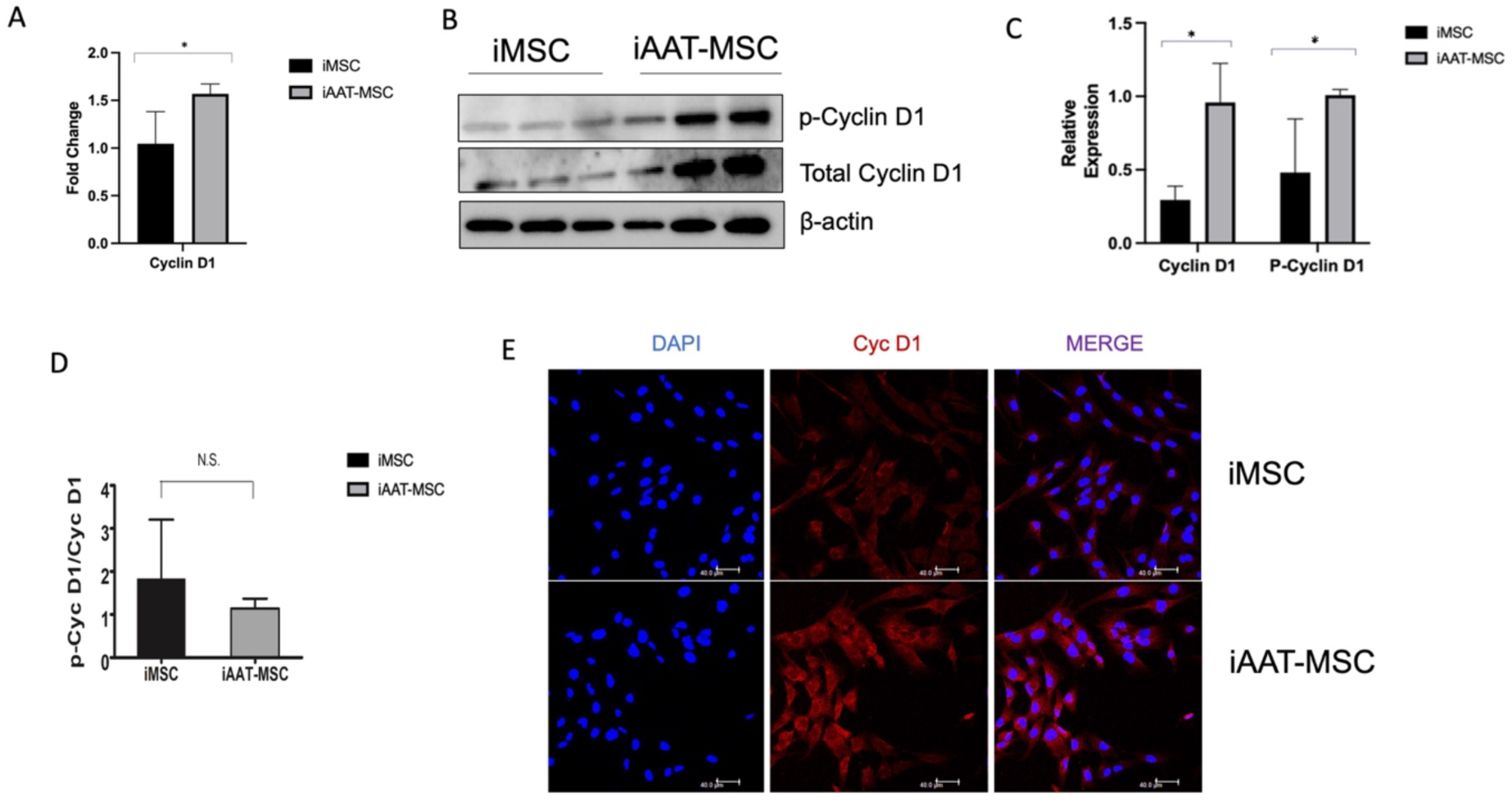
Cyclin D1 mRNA and protein expression in iAAT-MSCs and iMSCs. (A). Cyclin D1 mRNA expression was higher in iAAT-MSCs compared to iMSCs (*p<0.05); data are fold changes compared to GAPDH expression in each sample. (B) Expression of Cyclin D1 and phosphorylated (p)-Cyclin D1 and β-actin in iMSCs and iAAT-MSC. Lanes 1-3 are iMSCs in triplicate and 4-6 are iAAT-MSCs in triplicate. (C). Quantification/relative expression of cylinD1/β-actin and p-Cyclin D1/β-actin, p-Cyclin D1/ total Cyclin D1 at the protein level and ratios (D) in iAAT-MSCs compared to iMSCs. * (p<0.05, NS: not significant). E. Representative expression of Cyclin D1 stained from cell culture of iMSCs and iAAT-MSCs. Red signals identify Cyclin D and blue cells are nuclei. Scale bar = 40 μm. *p<0.05, ** p<0.01, ANOVA.

### 4. Inhibition of Cyclin D1 with imperatorin reduces AAT’s pro-proliferation Effect

Imperatorin is a phytochemical that has been proven to inhibit Cyclin D1 expression and arrest cells in the G1 phase of the cell cycle (16). To assess whether Cyclin D1 is in part responsible for the increased cell proliferation in iAAT-MSCs, passage 7 iAAT-MSCs were treated with imperatorin at 25 or 125mM. Cells were then fixed for cell cycle analysis using flow cytometry. Non-treated iMSC and iAAT-MSCs were used as controls. Our data showed that treatment with imperator significantly restored the percentage of iAAT-MSCs in the G0/G1. Consequently, a significant decrease in the percentages of cells in both the S and G2/M phases were observed in iAAT-MSCs treated with imperator compared to non-treated iAAT-MSCs (p<0.05, **Figure 4A&B**), suggesting the increased proliferation in iAAT-MSCs is Cyclin D1 dependent.

**Figure 4:**
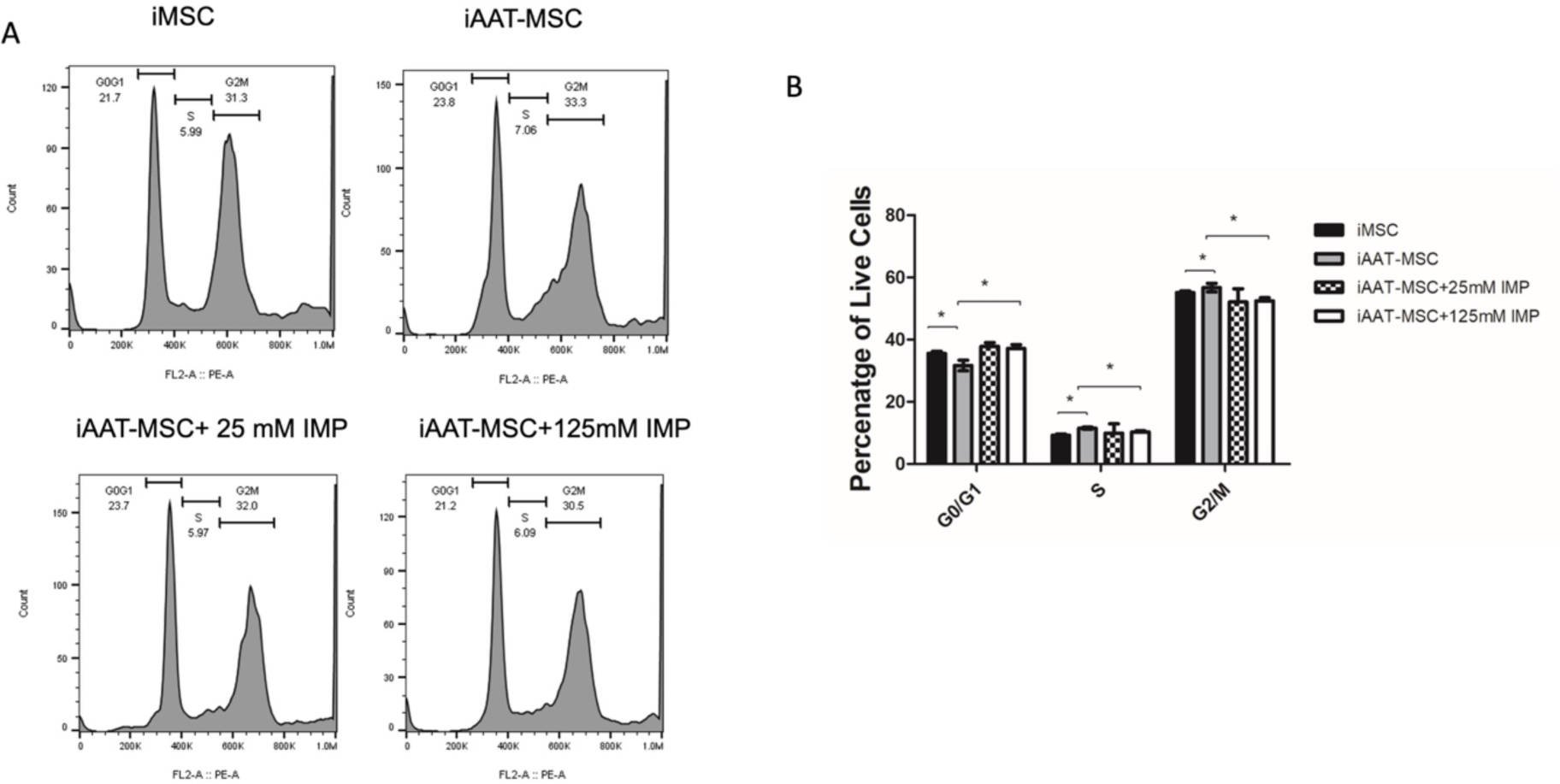
Inhibition of Cyclin D1 by imperatorin reduces cells in the S and G2/M stages in iAAT-MSCs. (A) Representative histogram of the cell cycle in iMSC, iAAT-MSCs and iAAT-MSCs treated with 25 or 125mM of imperatorin. IMP: imperatorin (B). Percentage of cells in each cell cycle phase in iMSC, iAAT-MSCs, and iAAT-MSCs treated with 25 or 125 mM imperatorin, p<0.05 by ANOVA test. *p<0.05, ANOVA. IMP: imperatorin.

### 5. Inhibition of Cyclin D1 with retinoic acid further confirms the role of Cyclin D1 in enhanced cell proliferation in iAAT-MSCs

Lipophilic retinoic acid (RA) inhibits Cyclin D1 activity by promoting ubiquitination and its proteolysis and arrests the cell cycle at the G1 phase (17). To confirm that the effect of AAT overexpressing on proliferation is Cyclin D1-dependent, we treated iAAT-MSCs with different 10 μM of RA, and then fixed the cells with PFA and performed cell cycle analysis. Again, iAAT-MSCs treated with RA had an increased percentage in the G0/G1 phase compared to non-treated iAAT-MSCs, and the level was comparable to iMSCs without any treatment (**Figure 5A&B**). This corresponded to a significant reduction in percentage of cells in the SA phase. These data again, suggest that increased iAAT-MSC proliferation was at least in part mediated by Cyclin D1.

**Figure 5:**
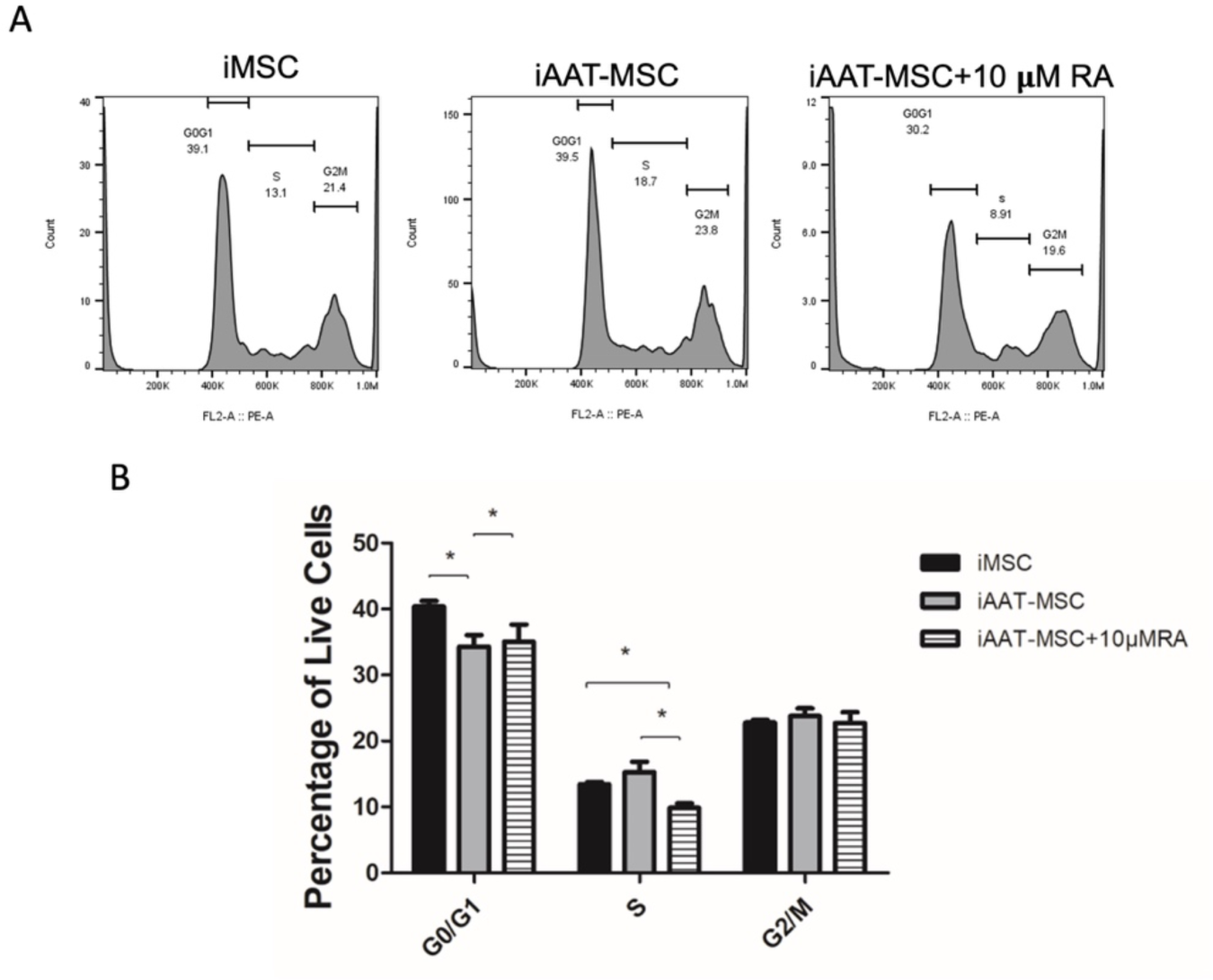
Retinoic acid inhibition of Cyclin D1 increases proliferation in iAAT-MSCs. (A) Representative histogram of cell cycle in iMSC, iAAT-MSCs and iAAT-MSCs treated with 10µM of retinoic acid. (B). Percentage of cells in each cell cycle phase in iMSC, iAAT-MSCs and iAAT-MSCs treated with retinoic acid, p<0.05 by ANOVA. Data shown are averages of at least three individual experiments. *p<0.05, ** p<0.01.

### 6. AAT-MSCs expressed a similar amount of GSK3β and β-Catenin compared to iMSCs

GSK3β, a serine/threonine protein kinase, plays a critical role in regulation of Cyclin D1 expression at both mRNA and protein levels (6). We next evaluated whether AAT overexpression led to enhanced GSK3β expression in iAAT-MSCs compared to iMSCs. Our Western blot data showed no significant differences in GSK3β or p-GSK3β expression between iAAT-MSCs and iMSCs (**Fig 6A-B**). There was also no difference in the ratio of p-GSK3β to total GSK3β between iMSCs and corresponding iAAT-MSCs (**Figure 6C**). Because the Cyclin D1 gene is also a target of the β-catenin pathway and the protein level of Cyclin D1 could be induced by β-catenin overexpression (18), we performed immunofluorescence staining in iMSCs and iAAT-MSCs for β-catenin expression. β-catenin signaling was observed in both the cytosol and inside the nuclei (activated form). There was no observable difference in the amount of activated (nucleus)/ cytosolic β-Catenin between iMSCs and iAAT-MSCs, suggesting that the upregulation of Cyclin D1 is likely independent of the Wnt signaling pathway.

**Figure 6:**
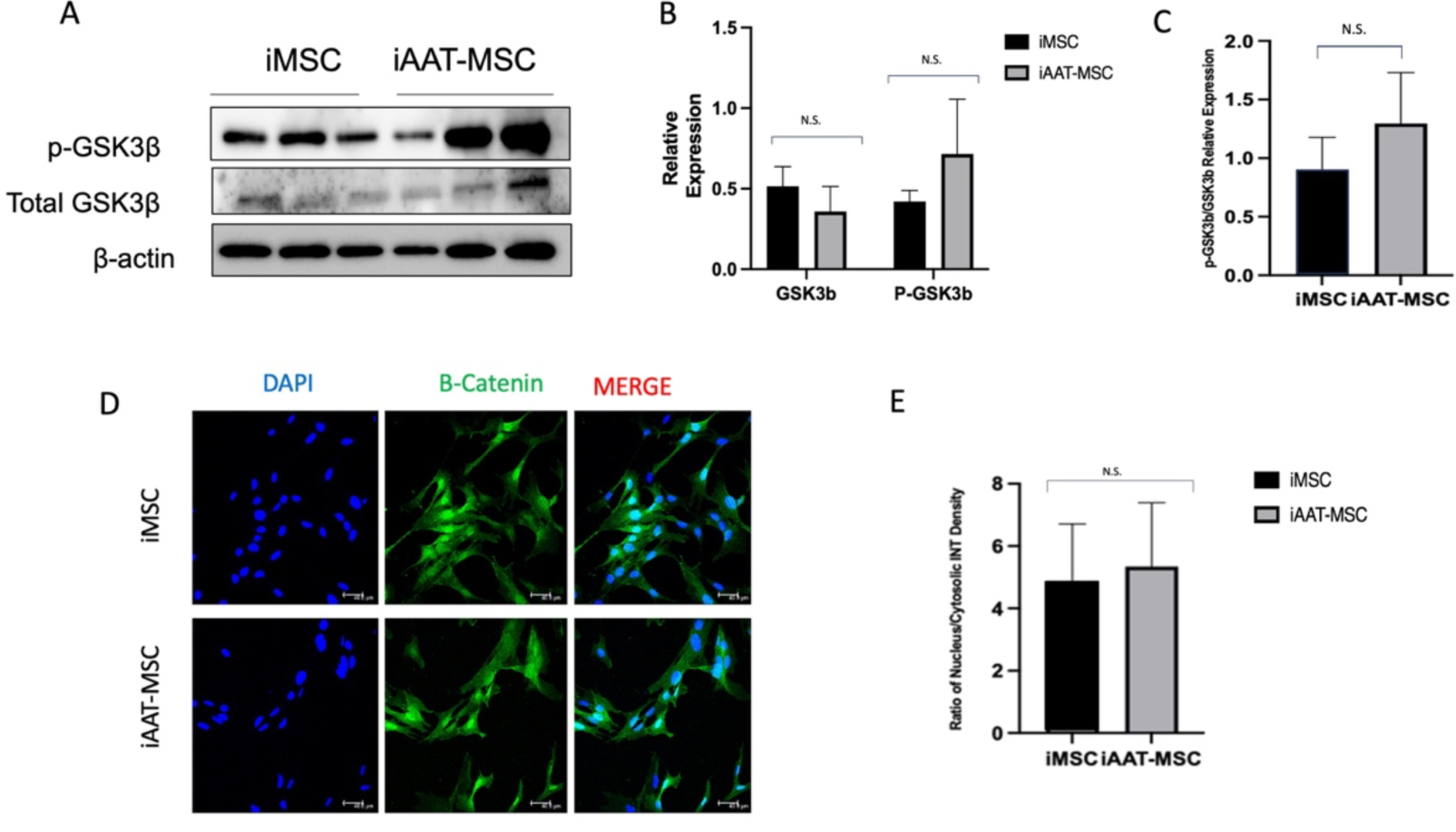
No change in expressions in GSK3β or β-catenin in iAAT-MSCs compared to iMSCs. (A). Western blot experiments performed in triplicate. Lanes 1-3 are iMSCs in triplicate and 4-6 are iAAT-MSCs in triplicate (same protein samples from Fig 3B, β-actin blot was used again for quantification). (B) Relative protein expression of GSK3β and p-GSK3β divided by β-actin, and (C) ratio of phosphorylated- and total GSK3β in iMSCs and iAAT-MSCs. (D) Immunofluorescence and (E) ratio of nucleus/cytosolic intensity of β-Catenin in iMSCs and iAAT-MSCs. NS: no significant difference.

## DISCUSSION

Mesenchymal Stromal/Stem Cells are being used therapeutically in a variety of diseases because of the potential for tissue specific responses. What is known about the science of these cells is that they have capacity to migrate throughout the body and a propensity to hone to areas of tissue injury. Exploring the signals associated with chemotaxis, migration, and cell to cell interactions remains in its infancy for these cells and could be different between subpopulations of MSCs.

Previous studies have shown that AAT enhances the therapeutic benefits of MSCs and aids in the protection of tissue and injury repair (1–3). Understanding the effects of AAT on MSC growth and gene expression may improve its clinical application in cellular therapy. This paper was stimulated by the observation that AAT markedly increased migration in MSC cell lines when given in direct culture or supplied as a gene therapy. The next set of experiments sought to determine if AAT increased properties of stemness and cell survival. What was found in this set of experiments is that MSCs move through the cell cycle more quickly by the upregulation of Cyclin D1 independent from the Wnt pathway. This increase in proliferative capacity does not prove that the cells are more or less potent in their functional therapeutic capacity. However, this feature may prove important with autologous MSC therapies in which a short time between collection of cells and therapeutic use is envisioned.

We consistently observed an increase in the rate of cell growth in primary and immortalized AAT-MSCs compared to iMSCs. This suggests that the overexpression of AAT contributed to increased cell proliferation. Indeed, iAAT-MSCs showed shorter G0/G1 phase with longer S phase. Shorter G0/G1 phase may predispose iMSCs to be especially reactive to differentiation signals, while a long S phase may allow the maintenance of a higher proportion of cells in the euchromatic rather than heterochromatic state (19, 20).

Previous single cell RNAseq comparing gene expression in MSCs and AAT-MSCs from 3 bone marrow donors indicated increases of cyclin D1. We therefore hypothesized that AAT overexpression might impact cell cycle in a Cyclin D-dependent manner. Since Cyclin D1 is one of the main cellular proteins that causes a cell to overcome the G1 checkpoint and progress into S phase it became a protein of interest to explore. Our results show that a significant increase in both gene and protein expression of Cyclin D1. In addition, the ratio of phosphorylated Cyclin D/ total cyclin D was not different between iAAT-MSCs and iMSCs meaning that the increased expression of total Cyclin D1 seen in iAAT-MSCs is due to an increase in activated Cyclin D1. To prove that this increase is the reason why we see the accelerated cell cycle progression we used two different known Cyclin D inhibitors: imperatorin and retinoic acid that can inhibit Cyclin D1 (17, 21). Both inhibitors effectively showed a significant decrease in the number of cells in S phase after flow cytometry, meaning that Cyclin D1 is the cause of this proliferation phenomenon we see in iAAT-MSCs and that AAT is an upstream positive influencer of Cyclin D1 expression.

One of the major pathways linked to Cyclin D1 expression is the Wnt signaling pathway (22). Our data suggests that AATs effect is independent of this major cellular pathway and independent of normal GSK3β negative regulation. Our results show no difference in the amount of phosphorylated or total GSK3β and no change in the activated β-catenin between the iAAT-MSCs and iMSCs. Therefore, these two proteins which are normally major upstream regulators of Cyclin D1 seem to not be affected by AAT. Moreover, since β-Catenin activation is the major endpoint of the Wnt pathway it is likely that AAT’s effect on Cyclin D1 expression is independent of the Wnt pathway. However, it is very possible that AAT’s effect on Cyclin D1 may be by influencing other cellular pathways and further research into the exact mechanism by which AAT increases Cyclin D1 is needed.

Some limitations of our study are present. We used cell lines from a limited donor pool. More diversity in the sample groups and replication of our findings by others is needed. Although these studies were also performed on cells that had not been immortalized, the immortalization procedure allowed us to use cells at later passage for many of these experiments. Whether immortalization impacted any of the proliferation signals remains unknown since the experiments were performed sequentially on native MSCs and native AAT-MSCs before replication in immortalized cell lines. However, we did not run these with concurrent controls. It is also worth noting that the only pathways our studies investigated were the Wnt signaling pathway and Cyclin D1. It is likely that AAT might also affect other cellular growth pathways and/or regulate the expression of other cyclin proteins.

In summary, our results indicate that AAT causes an increase in Cyclin D1 expression leading to an increase in proliferation in AAT overexpressing MSCs. Additionally, our results also suggest that this increased expression may be a direct effect of AAT independent of the Wnt signaling pathway.

## MATERIALS AND METHODS

### Cell preparation

MSCs were isolated from bone marrow specimens of three healthy donors purchased from the American Type Culture Collection (ATCC, Old Town Manassas, VA, USA). AAT-MSCs were prepared by lentiviral infection of native MSCs using a vector encoding human AAT gene with GFP as a reporter as reported previously (1). MSCs infected with control vectors were used as controls. MSCs and AAT-MSCs were expressing GFP and were sorted based on GFP expression. Immortalization was achieved by treating cells with simian virus 40 large antigen to generate immortalized MSCs (iMSCs) and immortalized AAT-MSCs (iAAT-MSCs) (23). Overexpression of AAT and presence of SV40 protein were confirmed by Western blot analysis (Shoebi, et al, unpublished data).

### Cell culture and cell counting

MSCs were cultured with Dulbecco’s modified Eagle’s medium (DMEM) supplemented with 10% fetal bovine serum, and 1% penicillin and streptomycin (Complete medium). For cell counting, cells were plated into 12 well plates at a seeding density of 2 x 10^5^ cells/well and allowed to grow in 5% CO_2_ at 37℃. Cells were then counted using a hemacytometer every 48 hours (h) for 7 days.

### Cell cycle analysis by Flow Cytometry

Cells were plated into 12 well plates (Corning) at a seeding density of 2 x 10^5^ cells and allowed to grow for 48 h in 5% CO_2_ at 37℃. Cells were collected, washed with PBS and centrifuged. The pellets were collected and fixed in 70% Ethanol at −20 ℃ overnight. After the fixation, cells were centrifuged at 1,000 RPM for 10 minutes. The supernatant was discarded, and the remaining pellets were washed and resuspended with a flow cytometry staining buffer (ThermoScience) and centrifuged at 1,500 RPM for 5 minutes. The pellets were resuspended in 500 μl of propidium iodide (PI) staining buffer and incubated for 15 mins at room temperature in the dark. Data of the stained living cells and their stage of development was then collected with a flow cytometer and further analyzed using FloJo software.

### Gene expression by qPCR analysis

Cells were plated into 12 well plates at a seeding density of 2 x 10^5^ cells and allowed to grow for 48 h in 5% CO_2_ at 37°C. Cells were then collected and suspended in a mixture of 350 μl RLT lysis buffer and 3.5 μl β-mercaptoethanol. RNA was extracted from this mixture using a RNeasy Micro Kit (Qiagen). The concentrations of RNA were measured using a BioTek Take 3 microplate reader. The concentrations were then standardized and reverse transcribed using the iScript cDNA Synthesis Kit to 500 ng/μl of cDNA. Advanced Universal SYBR Green Supermix (Bio-Rad) was used, and the quantitative RT-PCR was conducted in a CFX96 Real-Time Thermocycler (Bio-Rad). Fold changes in gene expression were normalized to GAPDH.

### Treatment of AAT-MSCs with Cyclin D inhibitors, imperatorin and retinoic acid

iAAT-MSCs were plated at a density of 5 x 10^5^ cells per well in 6-well plates and let to grow for 24 hours at 37°C. These cells were then treated with 25 mM or 125 mM of imperatorin, respectively. In a separate experiment, iAAT-MSCs were treated with 10µM retinoic acid. Cells were collected at 24 hours post-treatment for cell cycle analysis. Vehicle-treated iMSCs and iAAT-MSCs were used as controls.

### Western Blot

iMSCs and iAAT-MSCs were plated in 12 well plates at a density of 1 x 10^5^ cells/well and grew for 72 hours in a CO_2_ incubator at 37°C. The cells were then collected, washed with PBS and resuspended in a protein lysis buffer. Total protein was extracted, and protein concentration was measured using a BCA Protein Assay Kit (Thermo Fisher). For Western blot, 20 μg of protein was loaded and the proteins were separated by SDS-PAGE, transferred to PVDF membranes, and incubated with one of the following primary antibodies: goat anti-Cyclin D1/D2 (R&D Systems, cat: AF4196), rabbit anti-Phosphorylated Cyclin D-Thr286 (cell signaling, Cat: 3300T)), rat anti-GSK3 β (R&D Systems, MAB2506), rabbit p-GSK3 β (Cell Signaling, Cat; 5558T), and rabbit anti-β-Actin (Cell Signaling, Cat 7074). Secondary antibodies included horseradish peroxidase (HRP)-conjugated anti-goat, anti-rabbit, or anti-rat IgG (Cell Signaling). Signals were visualized using an ECL detection kit (Thermo Scientific). Relative Protein expression was quantified using the ImageJ Software (Bio-Rad).

### Immunofluorescence staining

The cells were plated into 12 well plates (Corning) with microscope slides at a seeding density of 2 x 10^5^ cells and allowed to grow for 48 h in 5% CO_2_ at 37℃. Cells were then fixed with 4% paraformaldehyde for 10 mins and permeabilized using 1% Triton. These cells were then incubated with primary antibodies: goat anti-Cyclin D1/D2 (R&D Systems, cat: AF4196), rabbit anti-β catenin (Cell Signaling Technology, Cat#:8480S), and followed by incubation with corresponding secondary fluorescent antibodies (Thermo Fisher). The cell nucleus was stained by 4’, 6-diamidine-2-phenylindole (DAPI) for 10 minutes. Slides were then mounted, observed and imaged using a Leica SP5 confocal microscope. Intensities of β catenin in the nuclei and cytosol were quantified by the ImageJ software.

### Statistical analysis

For statistical analysis, *Student’s t* test was used to compare the difference between two group’s means. Multiple comparisons were done with ANOVA. To get a proper p-value, each experiment was performed in primary MSCs and repeated in immortal MSCs by two people independently at least three times. Similar results were observed and only data done in iMSC and iAAT-MSCs are shown. Data presented in the graphs is the mean with standard deviation (SD). P-values <0.05 are accepted as significant.

## ACKNOWLEGEMENT

This study was supported in part by the National Institutes of Health (DK105183, DK 120394 and DK118529, and DK125464) and the Department of Veterans Affairs (VA-ORD BLR&D Merit I01BX004536).

## AUTHORS CONTRIBUTIONS

BW contributed to data acquisition, data analysis and drafted the manuscript. PM contributed to data acquisition and analysis. ML contributed to data acquisition and drafting of the manuscript. HWei, contributed to study design, data acquisition, and data analysis and proof reading. EG contributed to data acquisition. HW and CS provided study concept and design and manuscript writing. All authors read and approved the final manuscript.

